# Differential Sustained and Transient Temporal Processing Across Visual Streams

**DOI:** 10.1101/358473

**Authors:** Anthony Stigliani, Brianna Jeska, Kalanit Grill-Spector

**Affiliations:** Psychology Department, Stanford University, Stanford, CA, 94305, USA; Stanford Neurosciences Institute, Stanford University, Stanford, CA, 94305, USA

**Author notes:** Kalanit Grill-Spector, Department of Psychology, 450 Serra Mall, Stanford, CA, 94305.

**Keywords:** ventral stream, temporal encoding, category selectivity, fMRI, human visual system

## Abstract

How do high-level visual regions process the temporal aspects of our visual experience? While the temporal sensitivity of early visual cortex has been studied with fMRI in humans, temporal processing in high-level visual cortex is largely unknown. By modeling neural responses with millisecond precision in separate sustained and transient channels, and introducing a flexible encoding framework that captures differences in neural temporal integration time windows and response nonlinearities, we predict fMRI responses across visual cortex for stimuli ranging from 33 ms to 20 s. Using this innovative approach, we discovered that lateral category-selective regions respond to visual transients associated with stimulus onsets and offsets but not sustained visual information. Thus, lateral category-selective regions compute moment-to-moment visual transitions, but not stable features of the visual input. In contrast, ventral category-selective regions respond to both sustained and transient components of the visual input. Responses to sustained stimuli exhibit adaptation, whereas responses to transient stimuli are surprisingly larger for stimulus offsets than onsets. This large offset transient response may reflect a memory trace of the stimulus when it is no longer visible, whereas the onset transient response may reflect rapid processing of new items. Together, these findings reveal previously unconsidered, fundamental temporal mechanisms that distinguish visual streams in the human brain. Importantly, our results underscore the promise of modeling brain responses with millisecond precision to understand the underlying neural computations.

**AUTHOR SUMMARY:** How does the brain encode the timing of our visual experience? Using functional magnetic resonance imaging (fMRI) and a temporal encoding model with millisecond resolution, we discovered that visual regions in the lateral and ventral processing streams fundamentally differ in their temporal processing of the visual input. Regions in lateral temporal cortex process visual transients associated with stimulus onsets and offsets but not the unchanging aspects of the visual input. That is, they compute moment-to-moment changes in the visual input. In contrast, regions in ventral temporal cortex process both stable and transient components, with the former exhibiting adaptation. Surprisingly, in these ventral regions responses to stimulus offsets were larger than onsets. We suggest that the former may reflect a memory trace of the stimulus, when it is no longer visible, and the latter may reflect rapid processing of new items at stimulus onset. Together, these findings (i) reveal a fundamental temporal mechanism that distinguishes visual streams and (ii) highlight both the importance and utility of modeling brain responses with millisecond precision to understand the temporal dynamics of neural computations in the human brain.

## INTRODUCTION

How do high-level visual areas encode the temporal characteristics of our visual experience? The temporal sensitivity of early visual areas has been studied with electrophysiology in non-human primates (1)-4) and recently using fMRI in humans (5), 6). However, the nature of temporal processing in high-level visual regions remains a mystery for two main reasons. First, the temporal resolution of noninvasive fMRI measurements is in seconds (7), an order of magnitude longer than the timescale of neural processing, which is in the order of tens of milliseconds. Second, while fMRI responses are roughly linear for stimuli lasting 3-10 s (8), responses in visual cortex exhibit nonlinearities both for briefer stimuli, which generate stronger than expected responses (5), 6), 8)-13), as well as for longer stimuli, which get suppressed due to adaptation (14). Since the standard approach using a general linear model (GLM) to predict fMRI signals from the stimulus (8) is inadequate for modeling responses to such stimuli, the temporal processing characteristics of human high-level visual cortex have remained elusive [but see (12), 14)-17)].

We hypothesized that if nonlinearities are of neural (rather than BOLD) origin, a new approach that predicts fMRI responses by modeling neural nonlinearities can be applied to characterize temporal processing in high-level visual cortex. Different than the GLM, which predicts fMRI signals directly from the stimulus, the encoding approach first models neural responses to the stimulus and from them predicts fMRI responses. Recent studies show that accurately modeling neural responses to brief visual stimuli at millisecond resolution better predicts fMRI responses than the GLM (5), 6), 18). In particular, an encoding model with two temporal channels – one sustained and one transient – predicts fMRI responses in early and intermediate visual areas across a wide range of stimuli varying in duration from 33 ms to 30 s (5). The encoding approach also enables testing a variety of temporal models and quantifying which best predicts brain responses. By building generative models of neural computations, this approach offers a key framework that can provide insights into multiple facets of temporal processing including integration time windows (19)-21), channel contributions (5), 18), 22)-25), and the nature of response nonlinearities (5), 6), 9)-12), 18).

We considered three hypotheses regarding temporal processing in high-level visual cortex. One possibility is that temporal processing characteristics are similar across high-level visual regions but differ from those of earlier stages of the visual hierarchy. This hypothesis is based on results from animal electrophysiology showing longer latencies of responses in higher-level regions compared to primary visual cortex, V1 (1), as well as research in humans showing longer temporal receptive windows (19), 20) and integration times (21) in ventral temporal cortex (VTC) and lateral temporal cortex (LTC) compared to early visual areas. A second possibility is that temporal processing is uniform across high-level regions that process a shared category (e.g. face-selective regions in VTC and LTC) but differs across regions that process different categories (e.g. face‐vs. body-selective regions). This prediction is based on data showing differential responses to long-duration (21 s) images in face‐vs. place-selective regions in VTC (14), as well as differential response characteristics to fast (8 Hz) visual stimulation in body-selective regions vs. other category-selective regions (15). A third possibility is that temporal processing differs across ventral and lateral visual streams rather than across categories. A large body of literature has documented that regions in LTC along the superior temporal sulcus (STS) show heighted responses to biological motion compared to stationary stimuli and other types of motion (26)-33), unlike regions in VTC that may represent the static aspect of the stimulus (30), 31), 34). This predicts that lateral regions may show larger transient responses than ventral regions, which instead may show larger sustained responses.

To test these predictions, we measured fMRI responses in high-level visual areas to images of faces, bodies, and words that were either sustained (one continuous image per trial, durations ranging from 3–20 s) (**Fig. 1 *A-B***, *experiment 1*), transient [30 flashed, 33 ms long images per trial with interstimulus intervals (ISIs) ranging from 67–633 ms] (**Fig. 1 *A-B***, *experiment 2*), or contained both transient and sustained components (30 semi-continuous images per trial, durations ranging from 67–633 ms per image with 33-ms ISIs) (**Fig. 1 *A-B***, *experiment 3*). We also collected a separate functional localizer experiment to independently define regions selective to faces and bodies in VTC and LTC (**Fig. 1*C***; Materials and Methods). We used face‐and body-selective regions as a model system as there are multiple clusters of these regions across the temporal lobe, and face and body regions neighbor on the cortical sheet (35). This organization enabled us to (i) measure how the temporal dynamics of stimuli affect responses in each region and (ii) test if temporal processing characteristics vary across regions selective to different categories (e.g., faces or bodies) or across regions in different anatomical locations (e.g., ventral vs. lateral temporal cortex).

**Figure 1.**
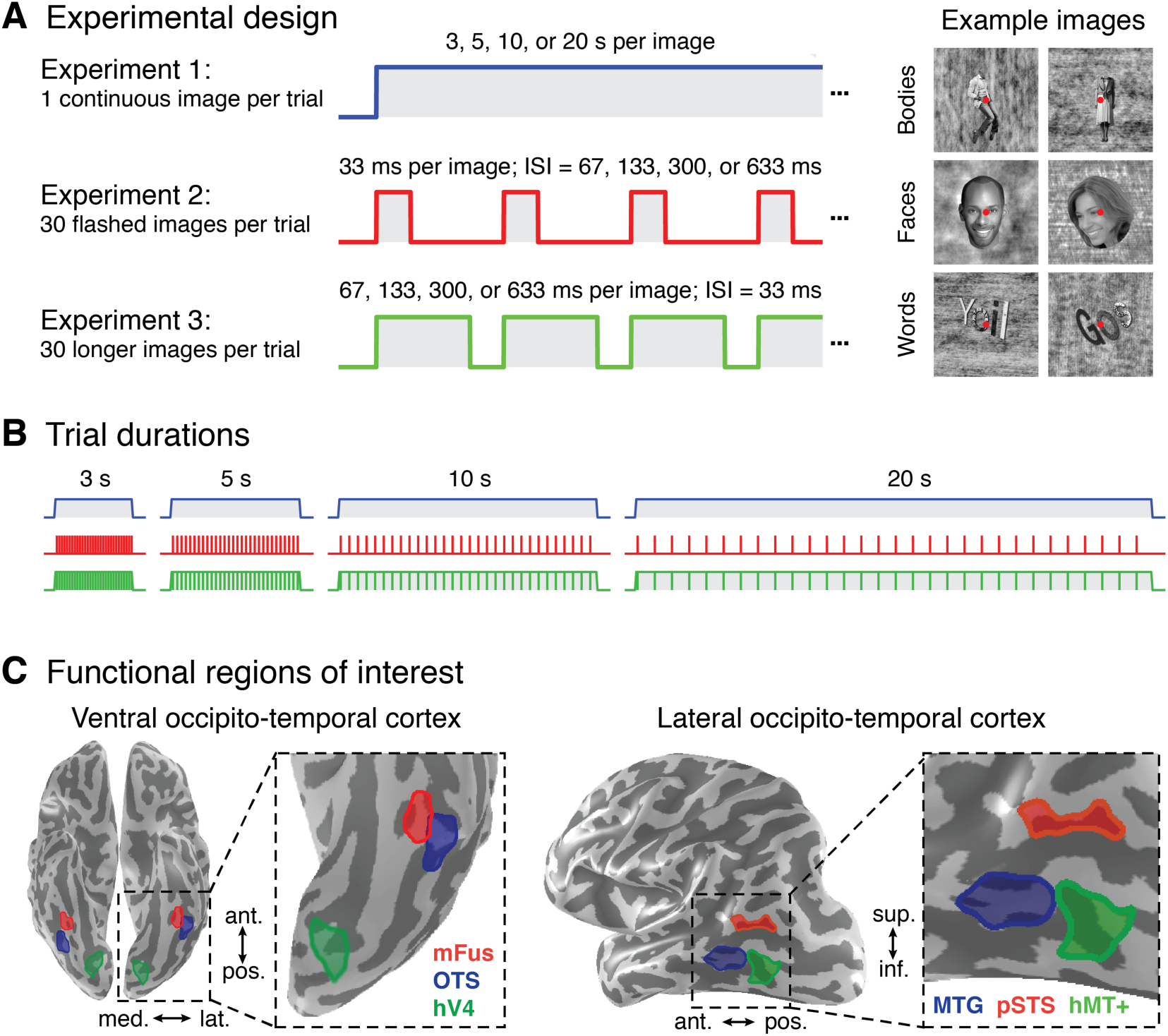
Measuring brain responses to combinations of sustained and transient visual stimuli in high-level visual cortex. (*A*) Participants fixated centrally and viewed images of bodies, faces, and pseudowords (*right*) that were presented in trials of different durations interleaved with 12-s periods of a blank screen (*left*). *Experiment 1:* a single image was shown for the duration of a trial. *Experiment 2:* 30 briefly presented images from the same category (33 ms each), each followed by a blank screen, were presented in each trial. As the trial duration lengthens, the gap between images increases, causing the fraction of the trial containing visual stimulation to decline. *Experiment 3:* 30 semi-continuous images from the same category were presented in each trial with a constant 33-ms blank screen between consecutive images. As the block duration lengthens, the duration of each image progressively increases but the gap does not. (*B*) The same trial durations (3, 5, 10, or 20 s) were utilized across all three experiments, while the rate and duration of visual presentation varied between experiments. Corresponding trials in experiments 1 and 3 have almost the same overall duration of stimulation but different numbers of stimuli, whereas trials in experiments 2 and 3 have the same number of stimuli but different durations of stimulation. The same fixation task was used in the three main experiments. (*C*) Functional regions of interest in ventral temporal cortex (*left*) and lateral temporal cortex (*right*) selective to bodies (OTS and MTG) and faces (IOG and mFus), as well as human V4 (hV4) and human motion-sensitive area (hMT+). Regions in each anatomical section are shown in an example subject.

## RESULTS

### Responses in High-Level Visual Cortex Exhibit Temporal Nonlinearities

To assess the feasibility of our approach, we first used a standard widely-used GLM (8) to predict fMRI responses in the three main experiments. Then, we compared these predictions to measured fMRI responses from two sample functional regions of interest: a ventral body-selective region and a lateral body-selective region.

In general, the GLM predicts longer responses for longer trials and similar responses in experiments 1 and 3 (**Fig. 2A**, *blue* and *green*). Responses in experiment 1 are predicted to be slightly higher than in experiment 3 because the 33-ms gaps between images in the latter experiment make up 1 s of baseline within each trial. Due to the nature of the hemodynamic response function (HRF), the GLM also predicts that peak response amplitudes in experiments 1 and 3 will increase gradually from 3-s to 10-s trials and subsequently plateau for longer trial durations. In contrast, this model predicts substantially lower responses in experiment 2 compared to the other experiments because the transient 33-ms stimuli in this experiment comprise only a small fraction of each trial duration (**Fig. 2*A***, *red*). Therefore, the GLM predicts a progressive decrease in response amplitude from 3-s to 20-s trials in experiment 2, as the fraction of the trial in which stimuli are presented decreases (from 1/3 to 1/20 of the trial).

**Figure 2.**
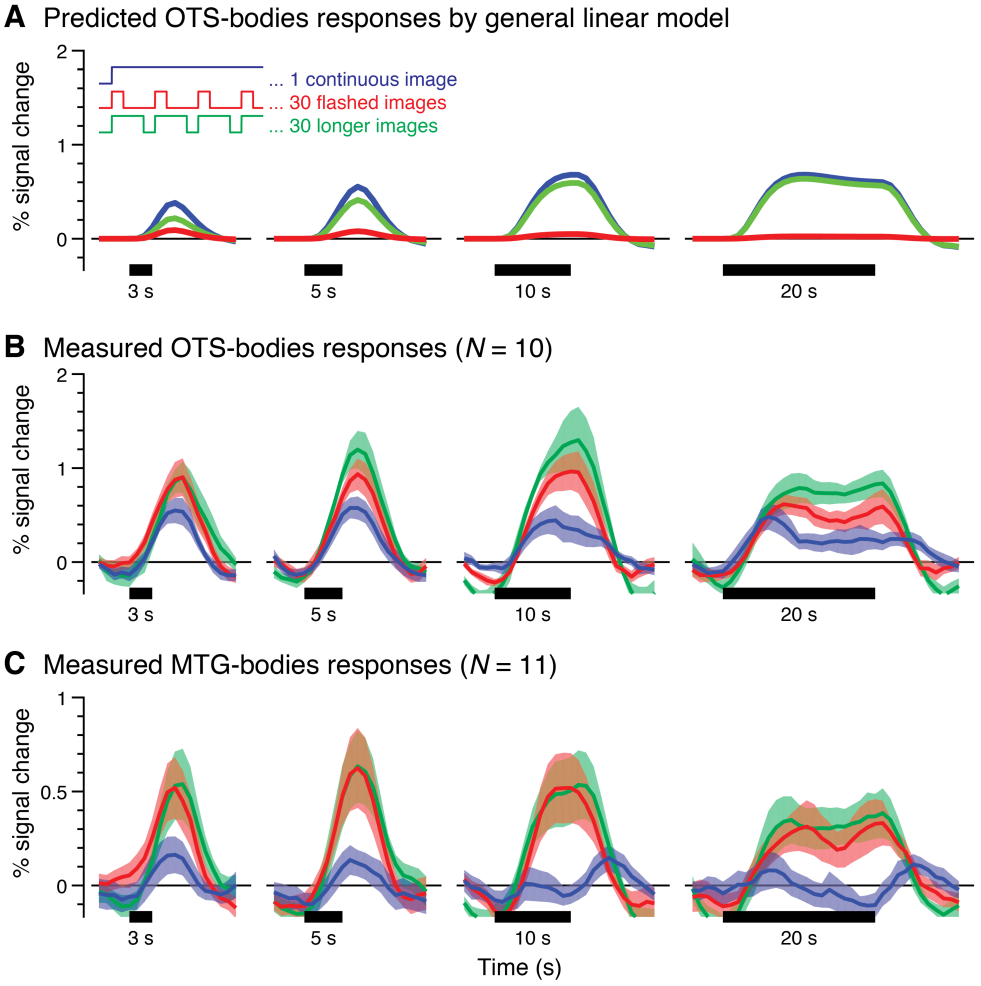
Responses of body-selective regions in ventral and lateral temporal cortex exhibit nonlinearities that are not predicted by a linear model. (*A*) Predicted responses by a GLM for trials containing one continuous image (*blue*), thirty flashed (33 ms) images (*red*), and thirty longer images that span then entire trial duration except for a 33 ms interstimulus interval (ISI) following each image (*green*). Predictors are fit to OTS-bodies responses using data concatenated across all three experiments shown in (B). (*B*) Measured responses in a ventral region on the occipitotemporal sulcus (OTS) selective to bodies (OTS-bodies) during the three experiments. (*C*) Measured responses in a lateral region on the middle temporal gyrus (MTG) selective for bodies (MTG-bodies) during the three experiments. In (B-C), *lines:* mean response time series across participants for trials with body images; *shaded areas:* standard error of the mean (SEM) across participants; *Horizontal black bars:* stimulus duration.

Strikingly, responses to body images in a ventral body-selective region (OTS-bodies; **Fig. 2*B***) and a lateral body selective region (MTG-bodies; **Fig. 2*C***) both deviate from the predictions of the GLM, but in different ways.

In contrast to the predictions of the GLM, responses in OTS-bodies to trials of 30 flashed images in experiment 2 (**Fig. 2*B***, *red*) are substantially higher than in corresponding trial durations in experiment 1, when one stimulus is shown per trial (**Fig. 2*B***, *blue*). This occurs despite the fact that stimuli are presented for only a small fraction of each trial duration in experiment 2 compared to experiment 1. Furthermore, peak response amplitudes do not increase with trial duration in experiment 1 as predicted the GLM. Instead, we observe a systematic decrease in response after the first few seconds of stimulation in the 10-s and 20-s trials, which is consistent with prior reports of fMRI adaptation for prolonged stimuli in nearby face‐and place-selective regions (14). Lastly, responses in experiment 3 (**Fig. 2*B***, *green*) exceed responses in both experiment 1 (which has only one image per trial but similar overall durations of stimulation) and experiment 2 (which has the same number of images per trial but shorter stimulus durations). This observation suggests that both the number of stimuli in a trial and their duration impact response amplitudes, as in earlier visual areas such as V1 and hV4 (**Fig. S1**).

Unlike OTS-bodies, MTG-bodies illustrates a largely transient response profile with substantially lower responses to the prolonged single images in experiment 1. Notably, for the 10-s and 20-s trials, we observe a transient response following both the onset and the offset of the image but no elevation of response in the middle of the trial (**Fig. 2*C***, *blue*). In contrast to the lack of robust responses in experiment 1, MTG-bodies shows surprisingly large responses to briefly flashed stimuli in experiment 2 (**Fig. 2*C***, *red*). Additionally, responses in MTG-bodies during experiment 2 (**Fig. 2*C***, *red*) and experiment 3 (**Fig. 2*C***, *green*), which have 30 of stimuli per trial but of different stimulus durations, are similar and both exceed responses in experiment 1 which has a single stimulus per trial. This suggests that, unlike ventral regions, stimulus duration has little impact on MTG-bodies responses, which resemble responses in neighboring motion-sensitive hMT+ (**Fig. S1**).

These data demonstrate that (i) varying the temporal characteristics of visual presentations in the millisecond range has a profound effect on fMRI responses in high-level visual cortex, (ii) the standard GLM is inadequate for predicting measured fMRI responses to these types of stimuli in high-level regions, in agreement with prior data in earlier visual areas (5, 6, 8-13), and (iii) even though OTS-bodies and MTG-bodies prefer the same stimulus category, their temporal response characteristics vastly differ.

### An Encoding Model of Temporal Processing in High-Level Visual Cortex

Motivated by the recent success of encoding models that predict fMRI responses in earlier visual areas by modeling neural temporal nonlinearities (5), 6), 18), we applied a similar approach to predict responses in high-level visual areas. Different than the GLM, the encoding approach first models the neural response in millisecond resolution and then convolves the estimated neural response with an HRF to predict fMRI responses (**Fig. 3**).

Our encoding model consists of two temporal channels (5), 18) – a sustained channel and a transient channel – each of which can be modeled using a neural temporal impulse response function (IRF) (2), 3), 5), 18), 36) followed by a nonlinearity. The sustained channel is modeled with a monophasic IRF (**Fig. 3*B***, *blue channel IRF*), which predicts a sustained neural response for the duration of the stimulus. To capture the gradual decay (adaptation, A) of response observed in ventral regions for sustained images (**Fig. 2*B***, *blue*), we apply a nonlinearity to the sustained channel in form of an exponential decay function (Materials and Methods). The transient channel is characterized by a biphasic IRF (**Fig. 3*B***, *red channel IRF*) that identifies changes to the visual input. That is, it acts like a derivative function, predicting no further increase in the neural response once a stimulus has been presented for longer than the duration of the IRF (5), 18). This channel too has a nonlinearity, as we hypothesize an increase in neural response at both the appearance (onset) and disappearance (offset) of a stimulus. To account for the pronounced transient responses in high-level visual regions (**Fig. S1**), we apply a flexible compressive nonlinearity on the transient channel using a pair of sigmoid (S) functions, one for the onset and another for the offset (Materials and Methods). Thus, we refer to this two channel temporal encoding model as the A+S model. The predicted fMRI response is generated by convolving the neural response predictors for each channel with the HRF and summing the responses of the two temporal channels (**Fig. 3*C***). Since the HRF acts as a low-pass filter, predicted fMRI responses can be downsampled with minimal distortion to match the slower sampling rate of fMRI measurements.

**Figure 3.**
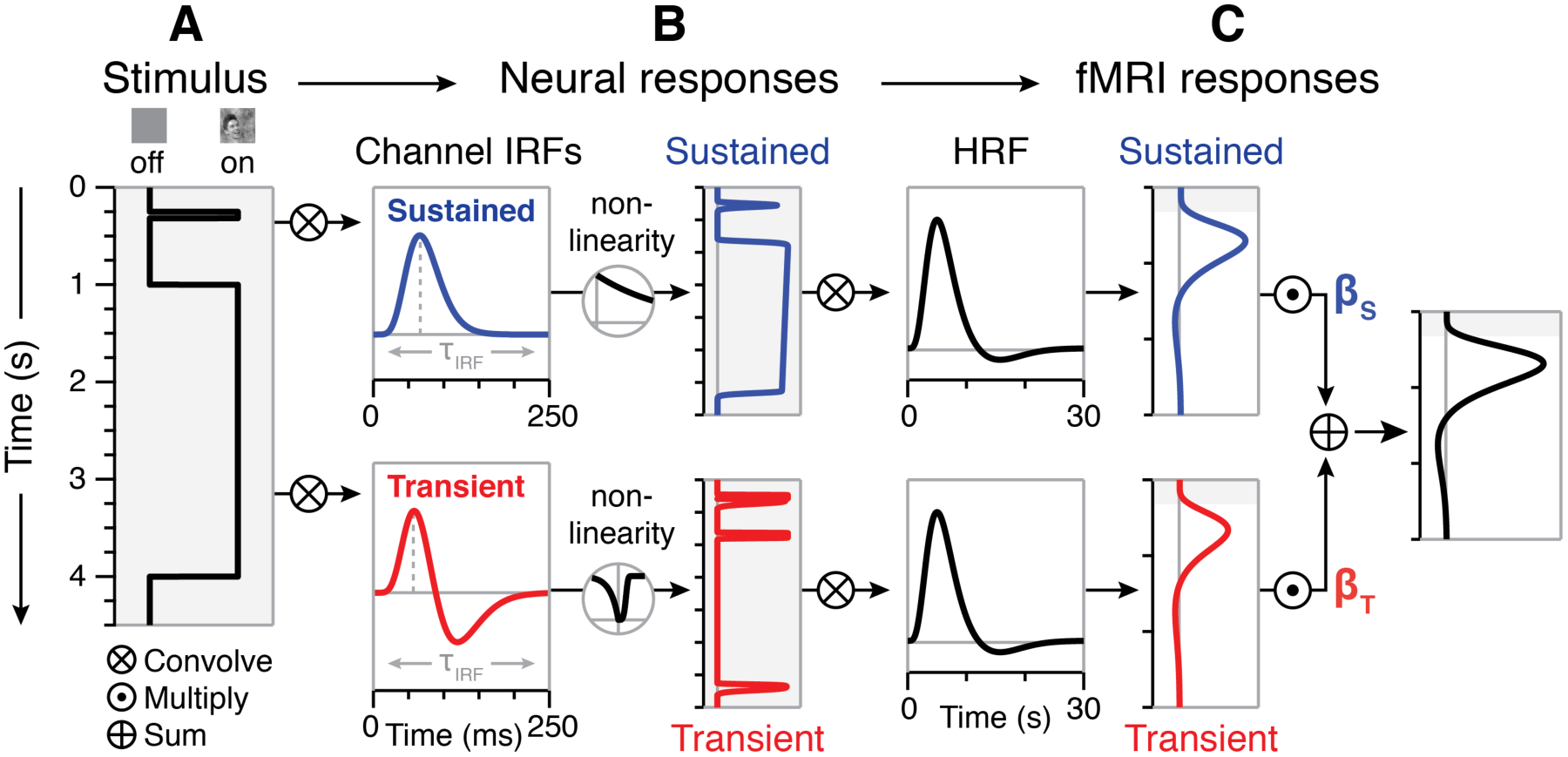
Optimized two-temporal channel A+S model with adaptation and sigmoid nonlinearities. (*A*) Transitions between stimulus and baseline screens are coded as a step function representing when a stimulus was on vs. off with millisecond temporal resolution. (*B*) Separate neural responses for the sustained (*blue*) and transient (*red*) channels are modeled by convolving the stimulus vector with an IRF for each channel. An exponential decay function is applied to the sustained channel to model response decrements related to neural adaptation, and a compressive sigmoid nonlinearity is applied to the transient channel to vary the temporal characteristics of “on” and “off” responses (Materials and Methods). (*C*) Predictors of sustained and transient fMRI responses are generated by convolving each channel’s neural response predictors with the HRF and down-sampling to match the sampling rate of measured fMRI data. The total fMRI response is the sum of the weighted sustained and transient fMRI predictors for each channel. To optimize model parameters and estimate the contributions (*β* weights) of the sustained (*β*_S_) and transient (*β*_T_) channels, we fit the model to different splits of the data including runs from all three experiments.

We estimated optimized A+S model parameters separately for each participant and region using nonlinear programming and a cross-validation approach. In our procedure, we use half the data from all three experiments to estimate model parameters. Specifically, we estimate a time constant for the neural IRFs (τ), a time constant controlling adaptation of sustained responses (α), and three parameters controlling compression of transient responses (*k*_on_, *k*_off_, and λ). After optimizing these parameters, we use a GLM to estimate the magnitude of response (*β* weight) for each channel and stimulus category in our experiments, resulting in three *β* weights for the sustained channel (one *β*_T_ for each category) and three *β* weights for the transient channel (one *β*_s_ for each category). These parameters and weights are then used to predict responses in left-out data and evaluate the model’s goodness-of-fit (cross-validated variance explained, x-*R*^2^).

Comparing the predictions of our optimized A+S model with measured fMRI responses in high-level visual cortex reveals two notable findings. First, our model generates signals that closely track the amplitude of fMRI responses in all three experiments in the left out data. Second, analysis of x-*R*^2^ shows that our optimized A+S model consistently outperforms other optimized temporal encoding models.

We illustrate these results for one region, OTS-bodies (**Figs. 4**, **S2**); Results for other regions are in **Figs. S3-S5**. Notably, the A+S model closely tracks response amplitudes in all three experiments [**Fig. 4 *A-C***, compare overall model prediction (*black*) with measured data from OTS-bodies (*gray*)]. Consistent with our predictions, the sustained channel accounts for the bulk of the response in experiment 1 (**Fig. 4*A***, *blue*); The transient channel contributes most of the response in experiment 2 (**Fig. 4*B***, *red*), and both channels contribute to responses in experiment 3 (**Fig. 4*C***).

**Figure 4.**
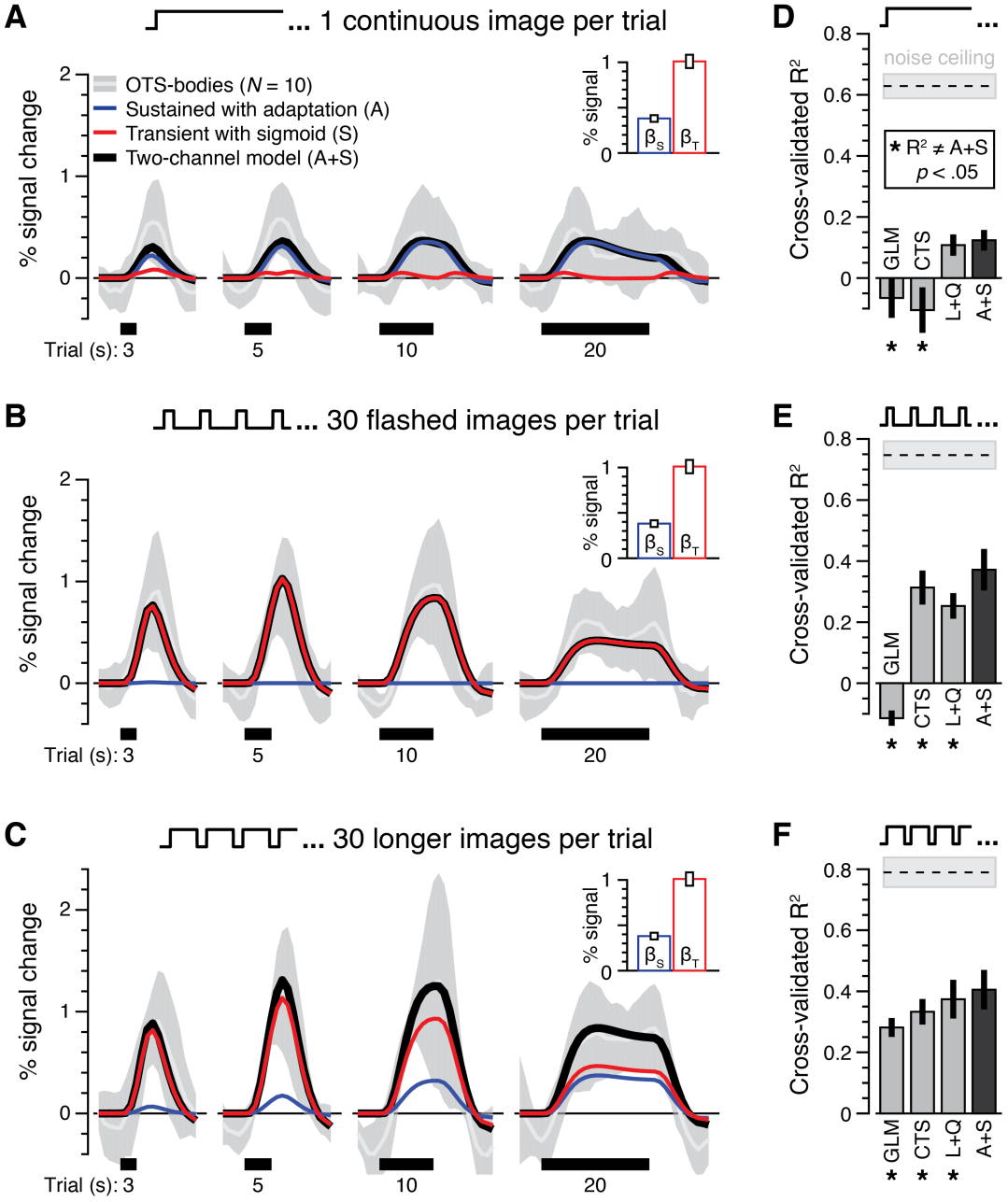
Two-temporal channel model with nonlinearities on both sustained and transient channels predicts responses in ventral temporal cortex. (A-C) Responses and model predictions for body images in OTS-bodies. *White curve:* mean response across 10 participants. *Shaded gray:* standard deviation across participants. *Blue:* predicted response from the sustained channel. *Red:* predicted response from the transient channel: *Black:* sum of responses from both channels. *Inset: mean* contribution (*β* weight) for each channel ±1 SEM across participants. (*A*) Experiment 1 data, 1 continuous image per trial. (*B*) Experiment 2 data, 30 flashed images per trial. (*C*) Experiment 3 data, 30 longer images per trial. (D-F) Model comparison. Bars show the performance of various models for each experiment presented in (A-C). Models are fit using runs from all three experiments, and cross-validation performance (x-*R*^2^) is calculated in left-out data from each experiment separately. (*D*) Experiment 1. (*E*) Experiment 2. (*F*) Experiment 3. Single-channel models: G*LM,* general linear model (8); *CTS,* a sustained channel with compressive temporal summation (6). Dual-channel models: *L+Q,* a linear sustained channel and a transient channel with quadratic nonlinearity (5); *A+S:* a sustained channel with adaptation and a transient channel with sigmoid nonlinearities. Asterisks denote models with significantly different performance compared to A+S (paired *t*-tests comparing x-*R*^2^ of each model vs. A+S in each experiment).

We compared the performance of our A+S model to other models of fMRI responses: the GLM (8), the balloon model (7), four single channel models (L, CTS (6), A, S; **Fig. S3*A***), and three alternative two-channel models (L+Q (5), 18), C+Q, A+Q) across all three experiments (Materials and Methods; **Figs. S3-S5**). For simplicity, **Fig. 4 *D-F*** compares performance in OTS-bodies for our model vs. three others: the standard GLM (8), a single-channel model with compressive temporal summation (CTS) (6), and a two-channel model composed of a linear sustained channel and a transient channel with a quadratic nonlinearity (L+Q) (5), 18). The latter two models have been recently used to model temporal dynamics of early and intermediate visual areas. Notably, the A+S model predicts OTS-bodies responses in left out data significantly better than the GLM (8), which overestimates responses in experiment 1 and underestimates responses in experiment 2 (GLM vs. A+S: all *t*s > 2.64, *P*s < .05, paired *t*-tests on x-*R*^2^ separately for each experiments) (**Fig. S2*A***). The A+S model also outperforms the recently proposed CTS model (6) that enhances early and late portions of the neural response to a stimulus (CTS vs. A+S: all *t*s > 2.60, *P*s < .05). While the CTS model performs considerably better than the GLM in experiment 2, it overestimates responses in experiment 1 with a single continuous images and underestimates responses in experiment 3 with 30 longer images per trial (**Fig. S2*B***). In experiments 2 and 3, we also observe a significant advantage of the A+S model compared to the two-temporal channel L+Q model (5), 18), which underestimates the large responses to transient stimuli in experiment 2 (**Fig. S2*C***) (L+Q vs. A+S: *t*s > 4.06, *P*s < .05; the difference fell short of significance for experiment 1, *t*_9_ = 1.98, *P* = .08).

Thus, an optimized two-temporal channel model with an adaptation nonlinearity in the sustained channel and compressive sigmoid nonlinearities in the transient channel predicts fMRI responses to visual stimuli ranging from milliseconds to seconds in high-level visual cortex with greater accuracy than alternative models.

### How do Channel Contributions Differ Across Ventral and Lateral Category-Selective Regions?

Examination of response time series (**Fig. S1**) and channel weights (**Fig. 5**) in body‐and face-selective regions in VTC and LTC reveals prominent differences across ventral and lateral temporal regions.

**Figure 5.**
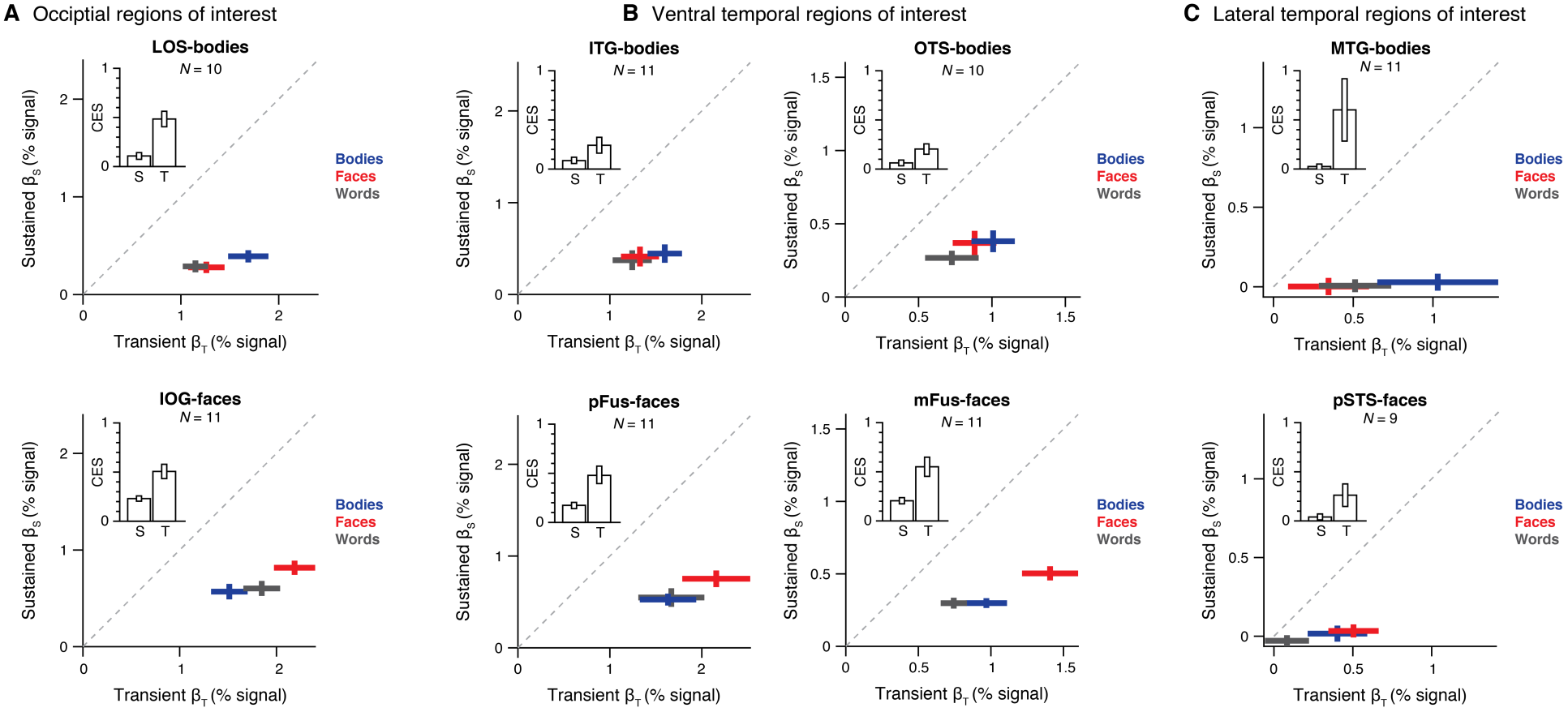
Differential contributions of transient and sustained temporal channels across ventral and lateral regions selective to face and body stimuli. Contributions (*β* weights) of transient (*x* axis) and sustained (*y* axis) channels for each stimulus category estimated by the two-temporal channel A+S model in (*A*) occipital body-selective region on the lateral occipital sulcus (LOS) and a face-selective region on the inferior occipital gyrus (IOGS), (*B*) ventral-temporal body-selective regions on the inferior temproal gyrus (ITG) and occipito-temporal sulcus (OTS)-bodies and face-selective regions on the posterior and mid fusiform gyrus, pFus‐and mFus-faces, respectivevely, and (*C*) a lateral temporal body-selective region on the mid temporal gyrus (MTG) and a face-selective region on the posterior aspect of the superior temporal sulcus (pSTS-faces). Crosses span ±1 SEM across participants in each axis, and *β* weights were solved by fitting the model using split halves of the data including runs from all three experiments. Data show average model weights across both splits of the data for each participant. *Red:* response to faces. *Blue:* response to bodies. *Gray:* response to words. *Dashed line*: identity line (*β*_S_ = *β*_T_). *Inset*: bars indicate smean contrast effect size (CES) of *β* weights for the preferred vs. nonpreferred categories in each channel ±1 SEM across participants.

First, comparing the response time courses of different category-selective regions shows that ventral temporal regions (e.g., OTS-bodies and mFus-faces) respond strongly to both the sustained stimuli in experiment 1 and the transient stimuli in experiment 2, whereas lateral temporal regions (MTG-bodies and pSTS-faces) respond strongly to the transient stimuli but minimally to the sustained stimuli (**Fig. S1**). The ratio of sustained and transient channel amplitudes, 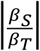 also differs across regions in ventral and lateral aspects of temporal cortex [significant main effect of processing stream, *F*_1, 107_ = 14.27, *P* < .01, three-way ANOVA with factors of processing stream (ventral/lateral), stimulus category (faces/bodies/words), and preferred category (bodies/faces)]. That is, while both sustained and transient channels contribute to responses in ventral temporal regions (**Fig. 5*B***), the transient channel dominates responses in lateral temporal regions (**Fig. 5*C***). In fact, zeroing the contribution of the sustained channel slightly improves model performance in lateral regions only (i.e. x-*R*^2^ of the S model is marginally better than the A+S model in MTG-bodies and pSTS-faces; **Fig. S3*B***). Moreover, ventral temporal regions show a characteristic similar to both hV4 (**Fig. S4*A***) and occipital category-selective regions (**Fig. 5*A***), whereas lateral temporal regions show a characteristic similar to motion-sensitive hMT+ (**Fig. S4*A***).

Second, in VTC (**Fig. 5*B***), category selectivity – or higher responses to a preferred category vs. other categories – is evident in both sustained and transient channels [all *t*s > 2.26, *P*s < .05, one-tailed *t*-tests comparing the contrast effect size (CES) of *β* weights for the preferred vs. nonpreferred categories separately for each channel; **Fig. 5*B***, *insets*]. For example, responses to faces in mFus are significantly higher than the average responses to words and bodies in both channels (*t*s > 5.25, *P*s < .001, paired *t*-test for each channel; **Fig. 5*A***, *right*), and responses to bodies in OTS are higher than average responses to other categories in both channels (*t*s > 2.59, *P*s < .05, paired *t*-test for each channel; **Fig. 5*A***, *left*). In contrast, in LTC (MTG-bodies and pSTS-faces; **Fig. 5*C***), there is a significant difference in the CES across sustained and transient channels [significant main effect of channel, *F*_1, 8_ = 14.88, *P* < .01, two-way ANOVA with factors of channel (sustained/transient) and preferred category (bodies/faces)]. That is, higher responses to the preferred category are observed only in the transient channel (*t*s > 1.99, *P*s < .05; the effect was not significant in the sustained channel, *t*s < 1.37, *P*s > .10; **Fig. 5*B***, *insets*). Thus, these results reveal differential contributions of transient and sustained channels across ventral and lateral category-selective regions. Finally, in the sustained channel, category-selectivity was higher in face-selective regions as compared to the body-selective regions.

### How do Timing Parameters Vary Across Ventral and Lateral Face and Body Regions?

We next examined the optimized timing and compression parameters for each channel across regions to test if there are functional differences across regions. The parameters in our A+S model were optimized separately for each region within each participant. Thus, for a given ROI, we optimized one time constant for the channel IRFs, one time constant for the adaptation decay function, as well as three sigmoid parameters (Materials and Methods).

For the sustained channel, we assessed how the time to peak of the neural IRF (*IRF*_S_) and the adaptation decay constant (α) vary across occipital and ventral temporal regions, omitting lateral temporal regions that did not have significant sustained responses. We discovered a hierarchical progression of longer time to peak and stronger adaptation in the sustained channel ascending from early to later stages of the ventral hierarchy (**Fig. 6*A***). That is, the time to peak of the sustained IRF tended to be shorter in V1 than hV4 and shorter in hV4 than in ventral regions OTS-bodies and mFus-faces (**Fig. 6*A***, *x axis*). A similar progression was observed from occipital to ventral temporal face‐and body-selective regions, whereby the time to peak of *IRF*_S_ for IOG-faces was shorter than mFus-faces and the time to peak of *IRF*_S_ for LOS-bodies was shorter than OTS-bodies. At the same time, the adaptation decay constant decreased from V1 to ventral temporal regions, indicating more adaptation in mFus-faces and OTS-bodies than in V1 (**Fig. 6*A***, *y axis*). Across face-selective regions we also observed a decreasing adaptation constant from IOG-faces/pFus-faces to mFus-faces, but our estimate of the adaptation constant was similar across occipital and ventral temporal body-selective regions. Thus, analyzing timing parameters in the sustained channel revealed hierarchical processing across the ventral stream, which was more salient in the estimate of the peak timing of the sustained *IRF*_S_ across regions.

**Figure 6.**
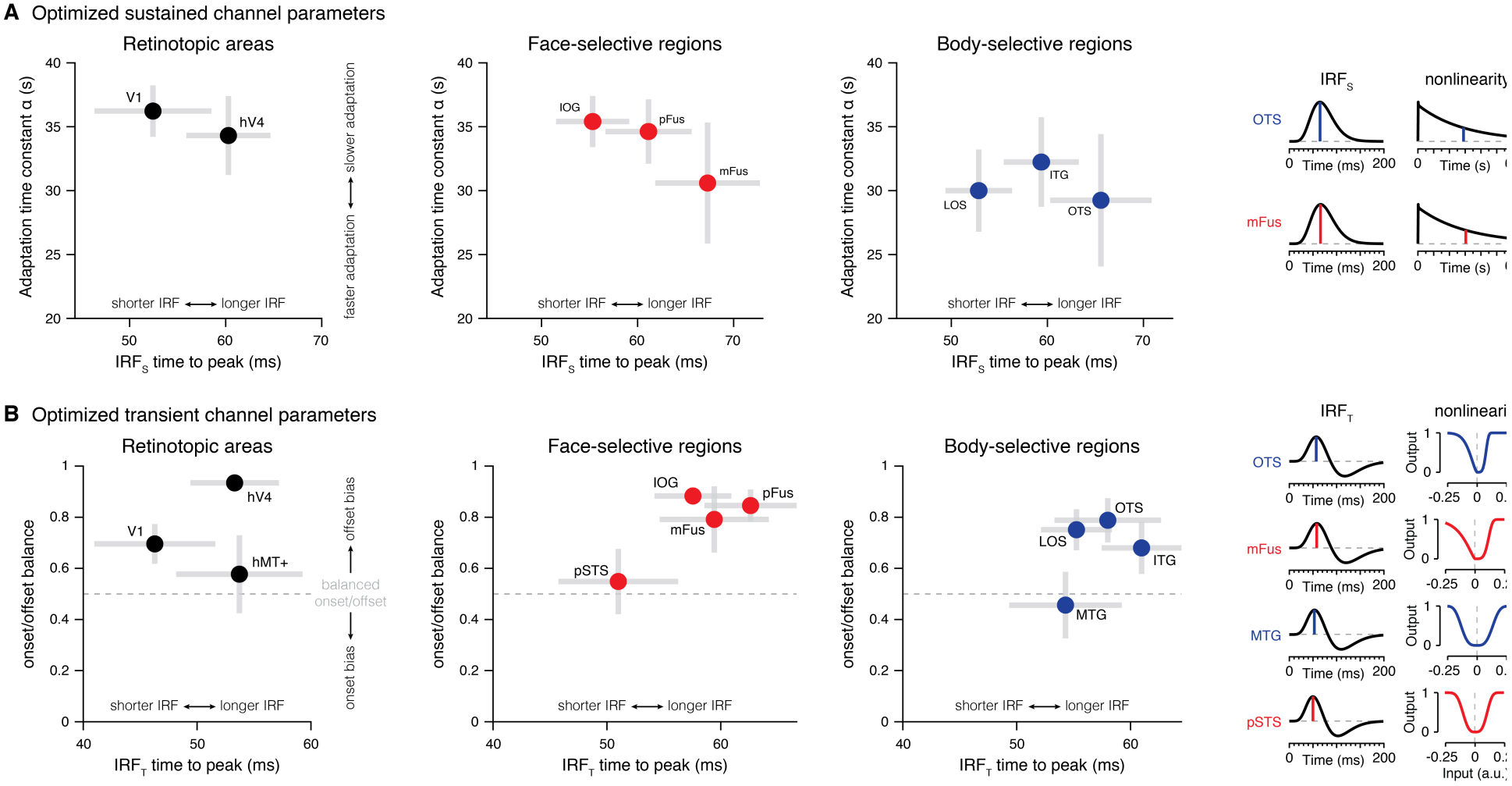
Optimized two-temporal channel model parameters differ across visual cortex. (*A*) Optimized sustained channel parameters. Time to peak of sustained *IRF*_S_ (*x* axis) and exponential time constant of the adaptation function (*y* axis) for each set of regions estimated by the two-temporal channel A+S model. Crosses span ±1 SEM across participants in each axis, and parameters were optimized using split halves of the data containing runs from all experiments. Data show model parameters averaged across both splits of the data for each participant. (*B*) Optimized transient channel parameters. Time to peak of transient *IRF*_T_ (*x* axis) and onset/offset balance (*y* axis) for each set of regions estimated by the two-temporal channel A+S model (with a zeroed sustained channel in lateral regions). The onset/offset balance metric captures differences in the shapes of the sigmoid nonlienarities used to compress transient “on” and “off” responses, where values larger than 0.5 refect elongation of offset responses compared to onset responses. Crosses span ±1 SEM across participants in each axis, and parameters were optimized using split halves of the data from all experiments. Plots show average model parameters across all splits of the data for each participant. Sample IRFs and nonlinearities shown to the right of (A-B) are generated by averaging optimized model parameters across participants.

In the transient channel, we examined how the time to peak of the *IRF*_T_ varies across regions and if there are asymmetries in the compression of “on” compared to “off” neural responses controlled by the sigmoid shape parameters *k*_on_ and *k*_off_, respectively. Since lower *k* values generally elongate transient responses, the relative contribution of the offset component can be indexed by a balance metric,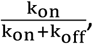 where a ratio of 0.5 indicates equal contributions from the onset and the offest of a stimulus to BOLD signals (*k*_on_=*k*_off_). A ratio < 0.5 indicates a larger contribution of onset than offset responses, and a ratio > 0.5 indicates a larger contribution of offset than onset responses.

First, like the sustained channel, the transient channel also shows an increase in the time to peak of *IRF*_T_ going from V1 to face‐and body-selective regions in VTC and LTC (**Fig. 6*B***, *x axis*). Second, VTC face-and body-selective regions tended to show longer time to peak of thier transient *IRF*_T_ as compared to LTC face-and body-selective regions. Third, interestingly, transients in lateral regions, pSTS-faces and MTG-bodies, show balanced contributions of onset and offset responses (balance metric = 0.50±0.09; **Fig. 6*B***, *y axis* and *insets*). In contrast, transients in ventral regions, pFus/mFus-faces and ITG/OTS-bodies, and occipital face-selective IOG and body-selective LOS) are dominated by offset responses (balance metric = 0.77±0.09; **Fig. 6*B*** and *insets*). The surprisingly large offset contribution in VTC indicates that the bulk of the response for the brief stimuli in experiment 2 can be attributed to neural responses that occur after the stimuli are no longer visible, rather than during the initial response to these stimuli.

Thus, comparison of optimized A+S model parameters reveals functional differences between early and later stages of the visual hierarchy, as well as distinct nonlinearities across ventral and lateral regions with the same category preference.

## DISCUSSION

Using a temporal encoding approach to explain responses in high-level visual regions, we discovered that an optimized two-temporal channel model consisting of a sustained channel with an adaptation nonlinearity and a transient channel with compressive sigmoid nonlinearities successfully predicts fMRI responses in human high-level visual cortex for stimuli presented for durations ranging from tens of milliseconds to tens of seconds. Critically, the innovative temporal encoding framework we introduce combines in a single computational model several components of temporal processing including time windows of temporal integration (12), 16), 19)-21), channel contributions (5), 18), 22)-25), and nonlinearities in temporal summation (5), 6), 9)-12), 18). Using this approach, we (i) uncover the temporal sensitivity of neural responses in human high-level visual cortex, (ii) find differential temporal characteristics across lateral and ventral category-selective regions, and (iii) propose a new mechanism – temporal processing – that functionally distinguishes visual processing streams in the human brain.

### Differences in Temporal Processing Across Visual Streams

Our results suggest two key differences between temporal processing in the ventral and lateral visual processing streams which project to ventral and lateral temporal cortex,respectively (37). First, there are differences in channel contributions. Lateral temporal cortex is dominated by responses to visual transients, while ventral temporal cortex responds to both sustained and transient visual information. Transient processing in LTC is consistent with the view that face and body-selective regions in the STS and MTG, respectively, are involved in processing dynamic visual information (26)-33). However, different than prior theories that have implicated these regions in processing biological motion (27)-29), 38), our data suggest that there is a more fundamental difference between high-level regions in lateral and ventral temporal cortex that is driven by differential channel contributions. Second, there are also differences in the dynamics of transient processing across visual streams. LTC regions show equal increases in neural responses due to the onset and offset of a visual stimulus, suggesting they carry information about moment-to-moment changes in the visual input. However, VTC regions exhibit surprisingly asymmetric contributions from the onset and offset of the stimulus. That is, the accumulation of fMRI responses due to the termination of a stimulus is more pronounced than responses associated with its onset. This difference suggests the intriguing possibility that transient responses in LTC code progressive changes to the visual input, while transient offset responses in VTC may reflect memory traces that are maintained in high-level regions after a stimulus is no longer visible. This prediction is consistent with results from ECoG studies showing that high frequency broadband responses (>60 Hz) in VTC continue for 100-200 ms after the stimulus is off (39)-42) and carry stimulus-specific information that may be modulated by attention (41), 42).

Observing a strong transient response in lateral regions, MTG-bodies and pSTS-faces, is interesting in the context of classic theories that propose differential contributions of magnocellular (M) and parvocellular (P) inputs to parallel visual streams in the primate visual system (22)-24). In macaques, the M pathway is thought to code transient visual information and projects from V1 to MT, while the P pathway is thought to code sustained information and projects from V1 to V4 and IT. While it is unknown how M and P pathways project to higher-level visual regions in the human brain, our results reveal that the transient channel dominates responses not just in hMT+ (5). This suggest that intriguing possibility that M projections not only dominate hMT+ as predicted by classic theories (22)-24), but also in surrounding face‐and body-selective regions in LTC.

Different from the predictions of classic theories of a predominant P input to the primate ventral stream (22)-24), we find significant contributions from both transient and sustained channels in VTC as well as evidence for category selectivity in VTC in both channels. This finding is consistent with later studies in macaques that reported that both M and P inputs propagate to ventral regions like V4 (5), 25). Surprisingly, our data in **Fig. 5** suggests that transient responses in VTC appear to be larger than sustained responses. We note that while interpreting the relative amplitude of responses within a channel is straightforward (e.g. comparing *β* weights for the different categories within the transient channel), interpreting the relative weight of sustained vs. temporal channels is complex, as it depends on the specific implementation of the model and the experimental design. Nonetheless, we are confident that there are both sustained and transient responses in VTC for two reasons. First, examination of raw responses during our experiments (**Fig. S1**), which are model free, shows that VTC regions respond strongly both to sustained single images (experiment 1) and trains of briefly flashed images (experiment 2). Second, responses in experiment 3, which had combinations of sustained and transient stimulation, exceed those of either experiment 1 or 2, suggesting additive contributions of the two channels.

Critically, finding substantial transient responses in VTC suggests a rethinking of the role of transient processing in the ventral stream. Specifically, it argues against the prevailing theoretical view that the ventral stream just processes static visual information. We hypothesize that transient responses in the ventral steam may serve two purposes. First, responses during onset transients may reflect rapid extraction of the gist of the visual input and may indicate the detection of a novel stimulus. Second, responses associated with offset transient, which were substantial in VTC may ignite a memory trace of the stimulus after it is no longer visible.

### What Are the Implications for Modeling fMRI Responses Beyond Visual Cortex?

Our data has critical implications for computational models of the brain. We developed a parsimonious yet powerful encoding model that can be applied to estimate nonlinear neural responses and temporal integration windows across cortex with millisecond resolution. While our two-temporal channel model provides a significant improvement in predicting fMRI signals compared to other models, we acknowledge that it does not explain the entire variance of the data. Future research may build upon the present results and improve model predictions by adding channels and nonlinearities. For example, examining adaptation effects on the transient channel as well as combining the temporal encoding approach with a spatial encoding approach (43)-45) may enable accurate prediction of brain responses to dynamic real-world scenes.

Given the pervasive use of the standard general linear model in fMRI research, our results have broad implications for fMRI studies of any part of the brain. We find that varying the timing of stimuli in the millisecond range has a substantial impact on the magnitude of fMRI responses. However, by estimating neural responses in millisecond resolution, we can accurately predict fMRI responses in second resolution for both brief and long visual stimuli. Thus, the temporal encoding approach we pioneered marks a transformative advancement in using fMRI to elucidate temporal processing in the brain because it links fMRI responses to the timescale of neural computations. As parallel streams occur not just in the visual system but throughout the brain, our data raise the intriguing hypothesis that temporal processing may also segregate other brain systems such as auditory or somatosensory cortex. Thus, our innovative approach offers a quantitative framework to identify functional and computational differences across cortex (46), 47) in many domains such as audition (48) and working memory (49). Importantly, the encoding approach can also be applied to study impairments in high-level abilities like reading (50) and mathematical processing (51) that require integrating visual information over space and time.

In sum, our results provide the first comprehensive computational model of temporal processing in high-level visual cortex. Our findings propose a fundamental new mechanism – temporal processing – that distinguishes visual processing streams whereby lateral category-selective process moment-to-moment visual transitions but ventral category-selective regions respond to both sustained and transient components. Visual transients in ventral category-selective regions may reflect rapid detection of changes to the visual content at stimulus onset and a memory trace of a recent stimulus at stimulus offset, which together suggest a new role of transient processing in the visual system beyond processing of dynamic stimuli. Finally, the encoding approach we introduce underscores the importance of modeling brain responses with millisecond precision to better understand the underlying neural computations.

## MATERIALS AND METHODS

### Participants

Twelve participants (6 males, 6 females) with normal or corrected-to-normal vision participated in the main experiments (experiments 1‒3). Each individual provided written informed consent and participated in two fMRI sessions: one session for experiments 1 and 2 and another session for experiment 3 and a functional localizer experiment (15). Seven participants from the main experiments (3 males, 4 females) also underwent population receptive field (pRF) mapping (43) to define retinotopic cortical regions and another experiment to define human motion-sensitive area (hMT+) (52). The Stanford Internal Review Board on Human Subjects Research approved all protocols.

### Temporal Channels Experiments

#### Visual stimuli

Stimuli consisted of well-controlled grayscale images of faces, bodies, and pseudowords (**Fig. 1 *A***, *right*) used in our previous publications (15). Stimuli were presented using an Eiki LCWUL100L projector (resolution: 1920 x 1200; refresh rate: 60 Hz) that was controlled by an Apple MacBook Pro using MATLAB (http://www.mathworks.com) and functions from Psychophysics Toolbox (53) (http://psychtoolbox.org). Participants viewed images through an auxiliary mirror mounted on the RF coil with stimuli spanning ∼20° of visual angle in each dimension.

#### Experimental design

To develop a temporal encoding model for high-level visual cortex, we adapted a fMRI paradigm previously used to model contributions of sustained and transient temporal channels in early visual cortex (5). The three main experiments in this study all used the same stimuli, trial durations, and task but varied the temporal presentation of the images. Critically, a 12-s baseline period (blank gray screen) always came before and after each trial. In all three experiments, participants were instructed to fixate on a small, central dot and respond by button press when it changed color (occurring randomly once every 2–14 s, 8 s on average).

*Experiment 1 ‒ one continuous image per trial:* Stimuli were shown in trials of varying durations (3, 5, 10, or 20 s per trial) in which a single image was shown for the entire trial. Acros trial durations the number of stimuli and transients (at the onset and offset of each stimulus) are matched but the duration of stimulation varies (**Fig. 1 *A-B***, *blue*). This experiment was designed enable measurement of fMRI-adaptation for prolonged images (14).

*Experiment 2 ‒ 30 flashed images per trial*: used the same trial durations as experiment 1, but in each trial we presented 30 different images from the same category. Each image was shown for 33 ms and followed by a blank interstimulus interval (ISI). Across trial durations the number of stimuli, number of transients, and total duration of visual stimulation are matched, but the ISI between consecutive images varied. Each ISI was 67 ms in the 3-s trials, 133 ms in the 5-s trials, 300 ms in the 10-s trials, and 633 ms in the 20-s trials (**Fig. 1 *A-B***, *red*).

*Experiment 3 ‒ 30 longer images per trial*: used the same design as experiment 2, except that in each trial we presented 30 images from the same category for longer durations with a constant ISI of 33 ms between images. Image durations varied across trials and were each shown for 67 ms in the 3-s trials, 133 ms in the 5-s trials, 300 ms in the 10-s trials, and 633 ms in the 20-s trials (**Fig. 1 *A-B***, *green*).

#### Data acquisition

Functional data were acquired using a simultaneous multi-slice EPI sequence with a multiplexing factor of 3 to obtain near whole-brain coverage with a TR of 1 s. Participants viewed four 270-s runs of each experiment. Each run of each experiment contained one instance of every permutation of stimulus category (face/body/word) and trial duration (3, 5, 10, or 20 s) presented in random order.

#### Category localizer experiment

To functionally define cortical regions that respond preferentially to specific stimulus categories, we collected three 300-s runs of a standard fMRI category localizer experiment used in our previous publications (15). Participants were instructed to fixate on a central dot and respond by button press when an image repeated randomly within a block. Code for the experiment is available at https://github.com/VPNL/fLoc.

#### pRF mapping and hMT+ localizer

To delineate retinotopic boundaries, we acquired four 200-s runs of pRF mapping (43) in a subset of participants from the main experiments. In this experiment, a bar swept across a circular aperture (40° × 40° of visual angle) in eight directions as participants performed a fixation task. To functionally define hMT+ in the same subset of participants, we collected one 300-s run of a fMRI motion localizer experiment as detailed in our prior publications (5), 52).

#### Magnetic resonance imaging (MRI)

MRI data were collected using a 3T GE Signa MR750 scanner at the Center for Cognitive and Neurobiological Imaging (CNI) at Stanford University.

*fMRI:* We used a Nova phase-array 32-channel head coil for the main experiments and functional localizer to obtain near whole-brain coverage (48 slices; resolution: 2.4 × 2.4 × 2.4 mm; one-shot T2*-sensitive gradient echo acquisition sequence: FOV = 192 mm, TE = 30 ms, TR = 1000 ms, and flip angle = 76°). We also collected T1-weighted inplane images to align each participant‘s functional data to their high-resolution whole brain anatomy.

For pRF mapping and the hMT+ localizer, we used a 16-channel visual array coil (28 slices; resolution: 2.4 × 2.4 × 2.4 mm; one-shot T2*-sensitive gradient echo acquisition sequence: FOV = 192 mm, TE = 30 ms, TR = 2000 ms, and flip angle = 77°) and collected T1-weighted inplane images in the same prescription.

*Anatomical MRI:* We acquired a whole-brain, anatomical volume in each participant using a Nova 32-channel head coil (resolution: 1 × 1 × 1 mm; T1-weighted BRAVO pulse sequence: FOV = 240 mm, TI = 450 ms, and flip angle = 12°).

### Data Analysis

Data were analyzed with MATLAB using code from vistasoft (http://github.com/vistalab) and FreeSurfer (http://freesurfer.net). Code used for predicting fMRI responses using a temporal channels approach is available at https://github.com/VPNL/TemporalChannels.

#### Region of interest (ROI) definition

Category-selective regions were defined in each participant’s native anatomical space at a common threshold (*t* > 3, voxel level, uncorrected) using functional and anatomical criteria detailed in prior publications (15) (**Fig. 1*C***). Face-selective ROIs (faces > others) were defined bilaterally in the inferior occipital gyrus (IOG-faces, *N* = 10), posterior STS (pSTS-faces, *N* = 9), posterior fusiform gyrus (pFus-faces, *N* = 11), and mid fusiform gyrus (mFus-faces; *N* = 11). Body-selective ROIs (bodies > others) were found bilaterally in the lateral occipital sulcus (LOS-bodies, *N* = 10), inferior temporal gyrus (ITG-bodies, *N* = 10), middle temporal gyrus (MTG-bodies, *N* = 11), and occipitotemporal sulcus (OTS, *N* = 10).

Visual areas V1 and hV4 were defined in each hemisphere in a subset of participant (*N* = 7) using data from the pRF mapping experiment. To match the visual field coverage of the stimuli in the main experiments, we restricted ROIs to only included voxels with pRF centers within the central 10°. We also defined bilateral hMT+ in the same subset of participants using data from the motion localizer experiment as in previous publications (5), 52).

#### Optimized two-temporal channel A+S model

To predict responses across all three experiments with a single model, we adapted an encoding approach introduced by prior studies (5), 18) that models fMRI responses as the weighted sum of activity across separate sustained and transient temporal channels.

In the procedure illustrated in **Fig. 3**, we first predict neural activity in each channel by convolving the stimulus time course in millisecond resolution (**Fig. 3*A***) separately with the neural IRF for the sustained channel (**Fig. 3*B***, *blue channel IRF*) and the transient channel (**Fig. 3*B***, *red channel IRF*). The sustained channel is characterized by a monophasic *IRF*_S_ that generates a response for the entire duration of a stimulus followed by an adaptation nonlinearity — convolved neural responses are multiplied by an exponential decay function beginning at the onset of each stimulus and extending until the onset of the following stimulus. In contrast, the transient channel is characterized by a biphasic *IRF*_T_ that generates a brief response at the onset and offset of an image (2), 3), 36). Here, convolved responses are passed through sigmoid nonlinearities that allow different levels of compression to be applied to the “on” and “off” responses. Then, the estimated neural responses for each channel are convolved with a hemodynamic response function (HRF) to generate a prediction of the fMRI response (**Fig. 3*C***). As such, there are neural nonlinearities in each channel of this model, but a linear relationship is assumed between the neural activity and BOLD responses. Finally, we use a GLM to solve for the contributions (*β* weights) of the sustained and transient channels, which reflect how much the predicted response from each channel is scaled before the responses of both channels are summed. Thus, the BOLD response to a stimulus can be expressed as

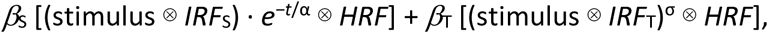

where *β*_S_ and *β*_T_ are the fitted response amplitudes for the sustained and transient channels, respectively; *IRF*_S_ and *IRF*_T_ are the impulse response functions for the sustained and transient channels, respectively; α determines the exponential decay at time *t* after stimulus onset; σ is a pointwise sigmoid nonlinearity, and *HRF* is the canonical hemodynamic response function.

#### Modeling nonlinearities in the neural response

We model the IRFs for each channel (**Fig. 3*B***) using formulas detailed in our prior publications (5). Here, we optimize the IRF time constant τ for each region, and the other parameters [taken from Watson (36)] are held constant: κ=1.33, *n*_1_=9, and *n*_2_=10.

Adaptation: To capture fMRI-adaptation (14) effects in the sustained channel, we use an exponential decay function, *e^−t^*^/α^, where *t* represents time after stimulus onset, and α indicates when the function declines to a proportion of 1/*e* (∼37%) of the initial response.

*Sigmoid nonlinearities:* To allow different levels of compression to be applied to “on” and “off” responses in the transient channel, we optimize separate sigmoid nonlinearities for the onset and offset responses using cumulative Weibull distribution functions,

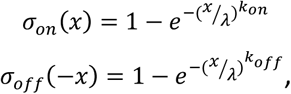

where λ is a sigmoid scale parameter used in both onset and offset nonlinearities; *k*_on_ is a sigmoid shape parameter that controls the curvature of the onset compression function, and *k*_off_ is a shape parameter controlling curvature of the offset compression function. Smaller *k* values produce more compressive nonlinearities that elongate transient “on” and “off” responses compared to larger *k* values.

#### Fitting and optimizing the two-temporal channel model

Since the HRF acts like a temporal low-pass filter, this allows resampling fMRI response predictors to the lower temporal resolution of the measured fMRI data (TR = 1 s) with minimal distortion. These resampled predictors are then compared with measured fMRI responses to estimate the contributions (*β* weights) of each channel for each category. To normalize the amplitude of predicted fMRI responses for the sustained and transient channels, we match the maximal height of predictors in the design matrix across the two channels. Finally, we used a GLM to estimate *β* weights of the sustained and transient channels for each stimulus category by comparing the predicted responses with the mean response time series of each ROI in each participant. To optimize the A+S model time constants (τ and α) and sigmoid parameters (λ, *k*_on_, *k*_off_) for each region, we used the constrained nonlinear optimization algorithm *fmincon* in MATLAB (*Optimization procedures*).

#### Validating the optimized two-temporal channel model

We assessed the predictive power of the optimized two-temporal A+S channel model by testing how well it predicts responses from separate runs of data from all three experiments. We first generated predicted neural response time courses by coding the visual stimulation in the left-out runs and convolving it separately with the IRFs of the sustained and transient channels (optimized using a separate split of the data). We then applied the adaptation and sigmoid nonlinearities, which were also optimized with independent data (**Fig. 6**). These transformed neural predictors were next convolved with the HRF and down-sampled to 1 s temporal resolution to match our fMRI acquisition. Finally, we multiplied each channel’s fMRI predictors with their respective *β* weights (estimated for each category in an independent split of the data) before summing the channel responses to predict fMRI responses. We then quantified how well the predicted responses matched the measured response across all data in the validation split.

Model performance was operationalized as cross-validated *R*^2^ (x-*R*^2^), which indexes the proportion of variance explained by *β* weights and model parameters that were estimated from independent data. While similar to a typical *R*^2^ statistic, x-*R*^2^ can be negative when the residual variance of a poor model prediction exceeds the measured variance in the response. Quantification of x-*R*^2^ within each experiment is presented in **Fig. 4 *D-F*** for OTS-bodies. Performance averaged across all three experiments is shown in **Figs. S3-S5** for all regions.

#### Testing alternative model architectures

To compare with the performance of our optimized two-temporal channel A+S model with alternatives, we tested five single-channel and three dual-channel models (**Figs. 2**, **4**, **S3-S5**).

*General linear model (GLM):* To first benchmark our model against a common GLM approach (8), we tested a linear model that predicts fMRI responses with a single convolution of the stimulus with the canonical HRF.

*Balloon model (B model):* To examine if responses in high-level visual cortex can be explained by a nonlinear hemodynamic model, we implemented the balloon model proposed by Buxton an colleagues (7) using standard parameters detailed in prior publications (5).

*Linear sustained channel (L model):* Similar to the GLM approach (8) but with two stages of convolution, we tested a single-channel model with a linear sustained channel,

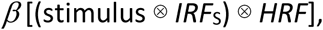

where *β* is a fitted response amplitude; *IRF*_S_ is the impulse response function for the sustained channel, and *HRF* is the canonical HRF.

*Sustained channel with compressive temporal summation (CTS*, **Figs. 4**, **S2-S5**): We also implemented a model proposed by Zhou et al. (6) composed of sustained channel with a compressive static power law,

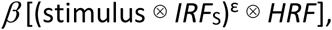

where ॉ is an optimized compression parameter ranging from 0–1.

*Sustained channel with adaptation (A model):* Identical to the sustained channel shown in **Fig. 3** (blue), we tested a model composed of a single sustained channel with adaptation,

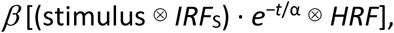

where α determines the exponential decay at *t* seconds after the onset of a stimulus.

*Transient channel with sigmoid nonlinearity (S model):* Identical to the transient channel shown in **Fig. 3** (red), we also tested a single-channel model composed of a transient channel with the same sigmoid nonlinearities described above,

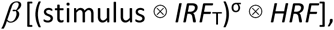

where *IRF*_T_ is the impulse response function for the transient channel and σ is a pointwise nonlinearity composed of separate sigmoid functions for onset and offset responses.

*Alternative dual-channel models (L+Q, C+Q, and A+Q models):* To compare the optimized two-temporal channel model shown in **Fig. 3** (A+S model) to alternative dual-channel models, we tested three variants of our model that all use a transient channel with a quadratic (Q) nonlinearity (squaring) but apply different nonlinearities in the sustained channel (**Fig. S3*A***). Combining different combinations of the sustained and transient channels described above, we compared two-channel models composed of a transient channel and either a linear sustained channel (L+Q model), a sustained channel with CTS (C+Q model), or a sustained channel with adaptation (A+Q model).

#### Optimization procedures

For all models with a neural IRF, we optimized a single time constant, τ, using formulas described in our prior publications (5). For models with adaption in the sustained channel (A and A+S), we also optimized an exponential time constant (α). For models with compressive temporal summation (CTS and C+Q models), we instead optimized an exponential compression parameter (ε). For models with a sigmoid nonlinearity in the transient channel (S and A+S models), we optimized three sigmoid parameters (λ, *k*_on_, *k*_off_). To optimize model parameters, we used the nonlinear optimization algorithm *fmincon* in MATLAB with the following constraints: τ = 4‒20 ms, α = 10‒40 s, ɛ = 0.01‒1, λ = 0.01‒0.5, *k*_on_ = 0.1‒6, and *k*_off_ = 0.1‒6 . The initial values passed to the optimizer for each parameter were τ = 4.93 ms, α = 20 s, ε = 0.1, λ = 0.1, *k*_on_ = 3, and *k*_off_ = 3. The cross-validation performance of each model averaged across all three experiments is shown in **Fig. S3 *B*** for category-selective regions in VTC and LTC and in **Figs. S4-S5** for other regions.

#### Statistical analyses

To test for differences in model performance across regions in VTC and LTC, we used a two-way repeated measures analysis of variance (ANOVA) with factors of model and region (comparing models and regions shown in **Fig. S3*B***). We then used paired two-tailed *t*-tests to compare the x-*R*^2^ of our model with others. **Fig. 4 *D-F*** contrasts the performance of our model (A+S) in OTS-bodies against three other models (GLM, CTS, L+Q) for each experiment individually. **Figs. S3-S5** contrast the performance of our model averaged across all three experiments vs. every other model for each region. To assess the level of noise in measurements from different brain regions, we also calculated a noise ceiling for each ROI using the inter-trial variability of responses for each condition as described in our prior publications (5). The noise ceiling estimate for OTS-bodies in each experiment is plotted in **Fig. 4 *D-F***, and the average noise ceiling across all three experiments is plotted in **Figs. S3-S5** for each region.

After establishing the validity of our model, we used paired two-tailed *t*-tests to compare β weights estimated by the A+S model for each region’s preferred category vs. average contributions for nonpreferred categories, separately for the sustained and transient channels. To test whether selectivity in the two channels differs across regions preferring bodies and faces in either VTC or LTC, we also used two-way ANOVAs with factors of channel (sustained/transient) and preferred category (bodies/faces) on the difference in channel weights for preferred vs. nonpreferred categories (contrast effect size, CES; **Fig. 5**). To examine whether the proportion of response attributed to sustained vs. transient channels differs across processing streams, stimulus categories, or regions preferring different categories, we then used a three-way ANOVA on channel contribution ratios, 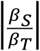 for each category with factors of stream (ventral/lateral), stimulus (faces/bodies/words), and preferred category (bodies/faces).

## ACKNOWLEDGEMENTS

We thank Jon Winawer and Jing Zhou for fruitful discussions. This research was supported by National Eye Institute Grant 1R01-EY02391501A1.

---

**Figure S1.**
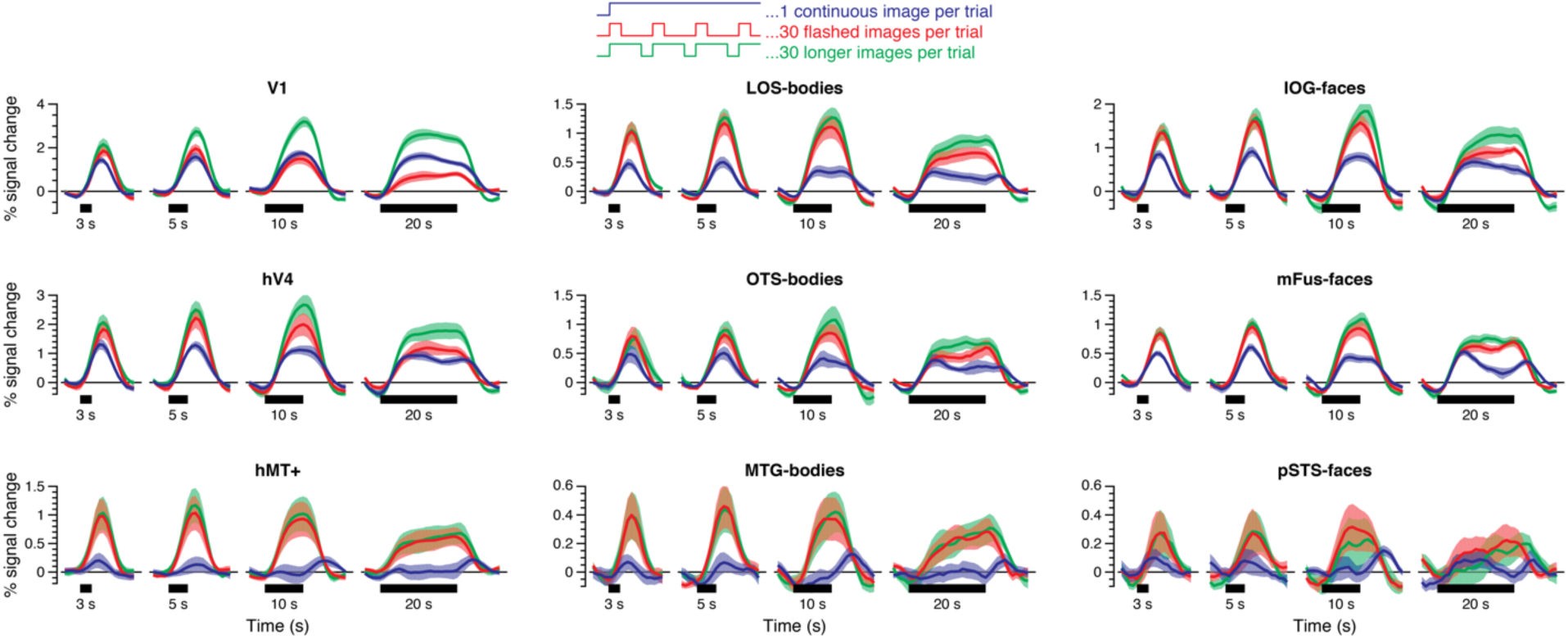
Responses to time-varying stimuli in occipital, ventral, and lateral regions of interest. Measured responses in occipital (V1, LOS-bodies, IOG-faces), ventral (hV4, OTS-bodies, mFus-faces), and lateral (hMT+, MTG-bodies, pSTS-faces) regions of interest in experiment 1 (*blue*), experiment 2 (*red*), and experiment 3 (*green*) averaged across all three stimulus categories. *Lines:* mean response time series across participants; *shaded areas:* standard error of the mean (SEM) across participants; *Horizontal black bars:* trial duration.

**Figure S2.**
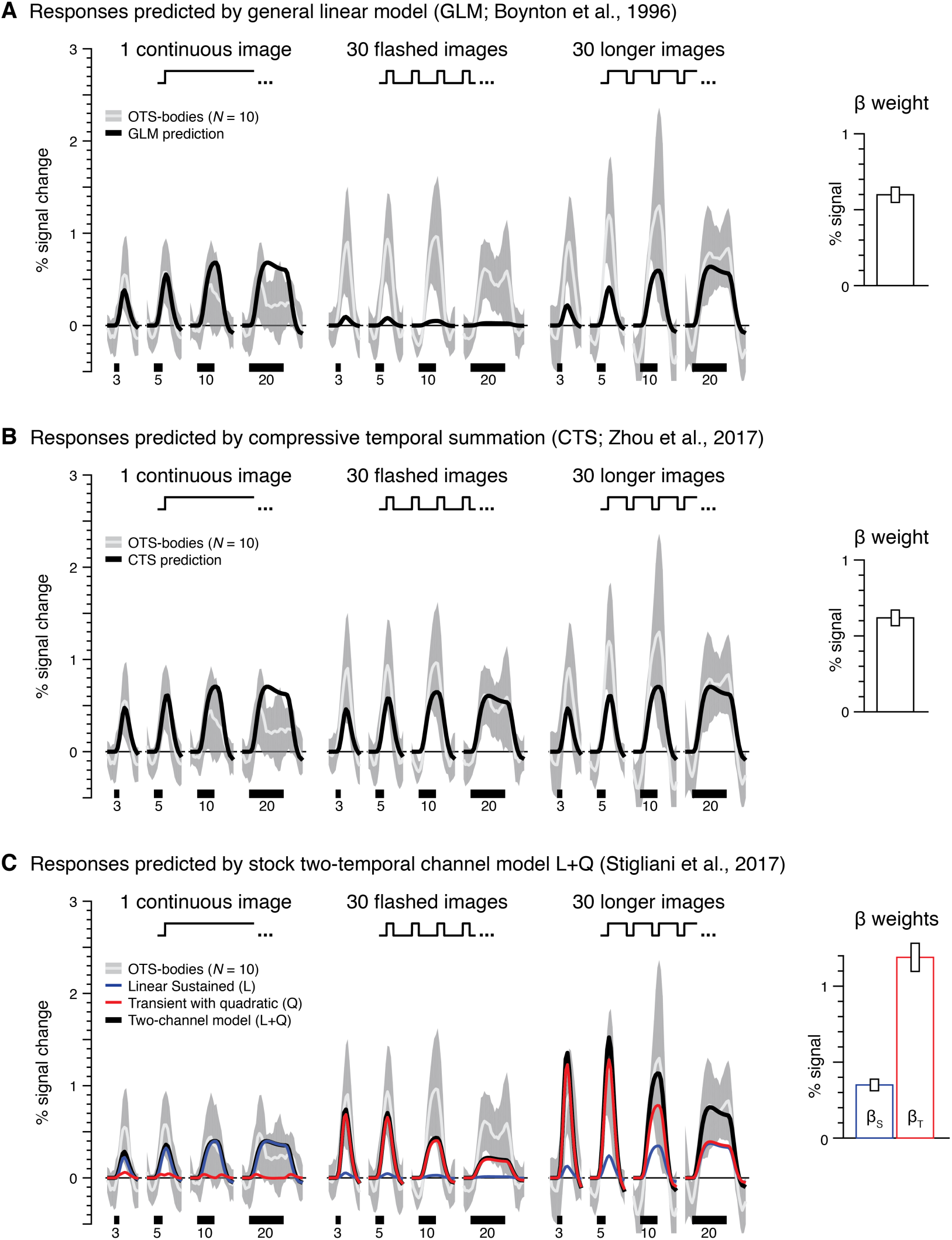
Comparison of temporal encoding models in OTS-bodies. (A-C) Responses and model predictions for body images in OTS-bodies for each experiment (*left*) with estimated *β* weights for each model (*right*). *White curve:* mean response across 10 participants. *Shaded gray:* standard deviation across participants. *Black curve:* overall model prediction. *Horizontal black bar:* trial duration. (*A*) Predictions of a general linear model (GLM) (8). (*B*) Predictions of a model with compressive temporal summation (CTS) (6). (*C*) Predictions of the two-temporal channel L+Q model with linear sustained channel and quadratic transient channel. *Blue curve:* predicted response from the sustained channel. *Red curve:* predicted response from the transient channel: *Black curve:* sum of responses from both channels.

**Figure S3.**
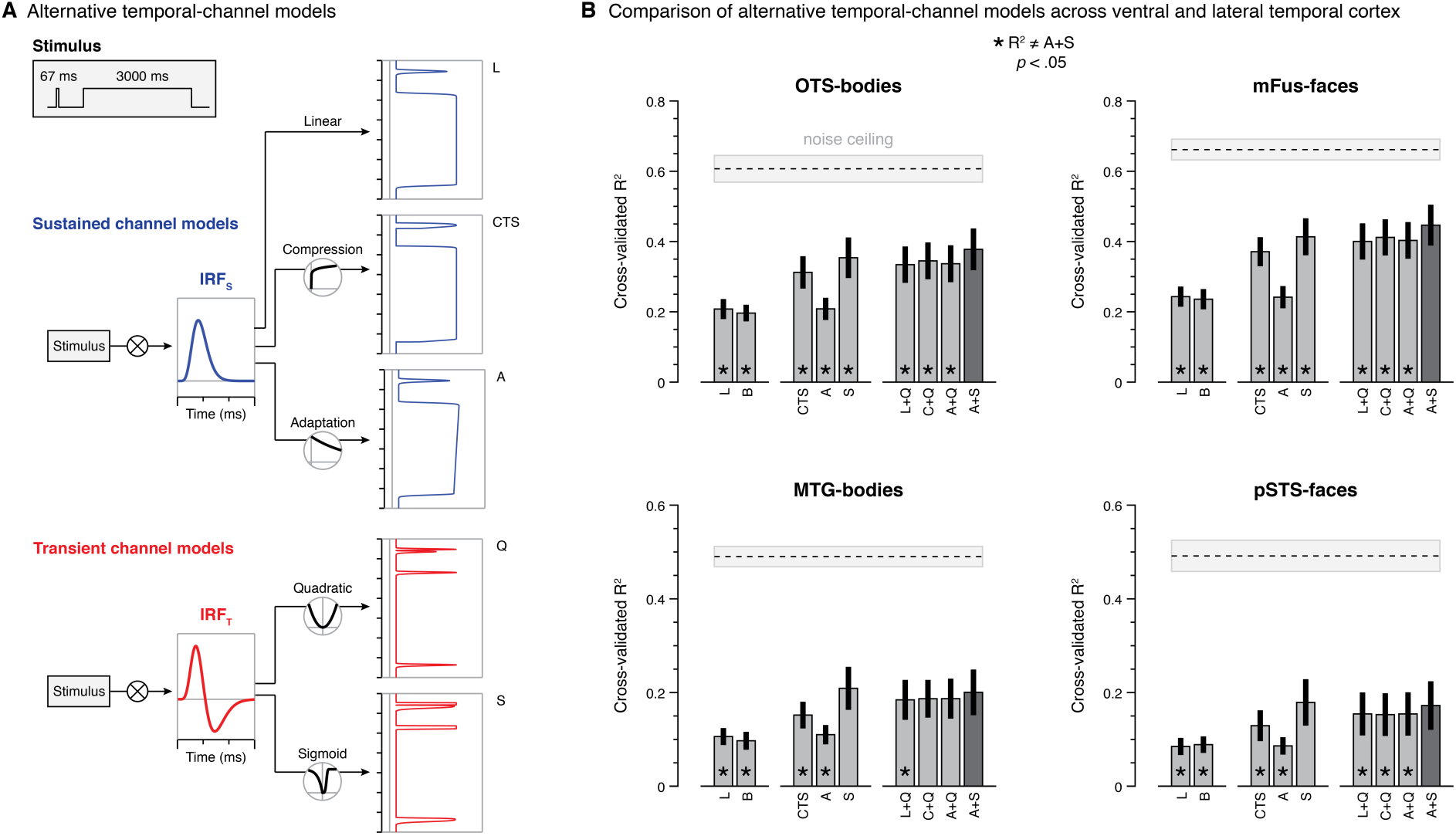
Comparison of temporal encoding models across visual cortex. (*A*) Alternative models of sustained (*blue*) and transient (*red*) channels. Schematic depicts neural response predictions generated by different implementations of each channel for both a brief (67 ms) and long (3 s) stimulus. Sustained channel models: *L,* a linear sustained channel; *CTS,* a sustained channel with compressive temporal summation (6); *A*, a sustained channel with adaptation (8). Transient channel models: *Q*, a transient channel with a quadratic (squaring) nonlinearity; *S*, a transient channel with a sigmoid nonlinearity. (*B*) Comparison of model performance (cross-validated *R*^2^) in each region averaged across all three experiments. Hemodynamic models: *L*, same as in (a); *B*, balloon model (7). Single-channel neural models: *CTS*, *A*, and *S*, same as in (a). Two-channel neural models: *L+Q,* a linear sustained channel and a transient channel with a quadratic nonlinearity (5); *C+Q,* a sustained channel with compressive temporal summation and a transient channel with a quadratic nonlinearity; *A+Q,* a sustained channel with adaptation and a transient channel with a quadratic nonlinearity; *A+S*, a sustained channel with adaptation and a transient channel with a sigmoid nonlinearity. Asterisks denote models with significantly different performance vs. the A+S model.

**Figure S4.**
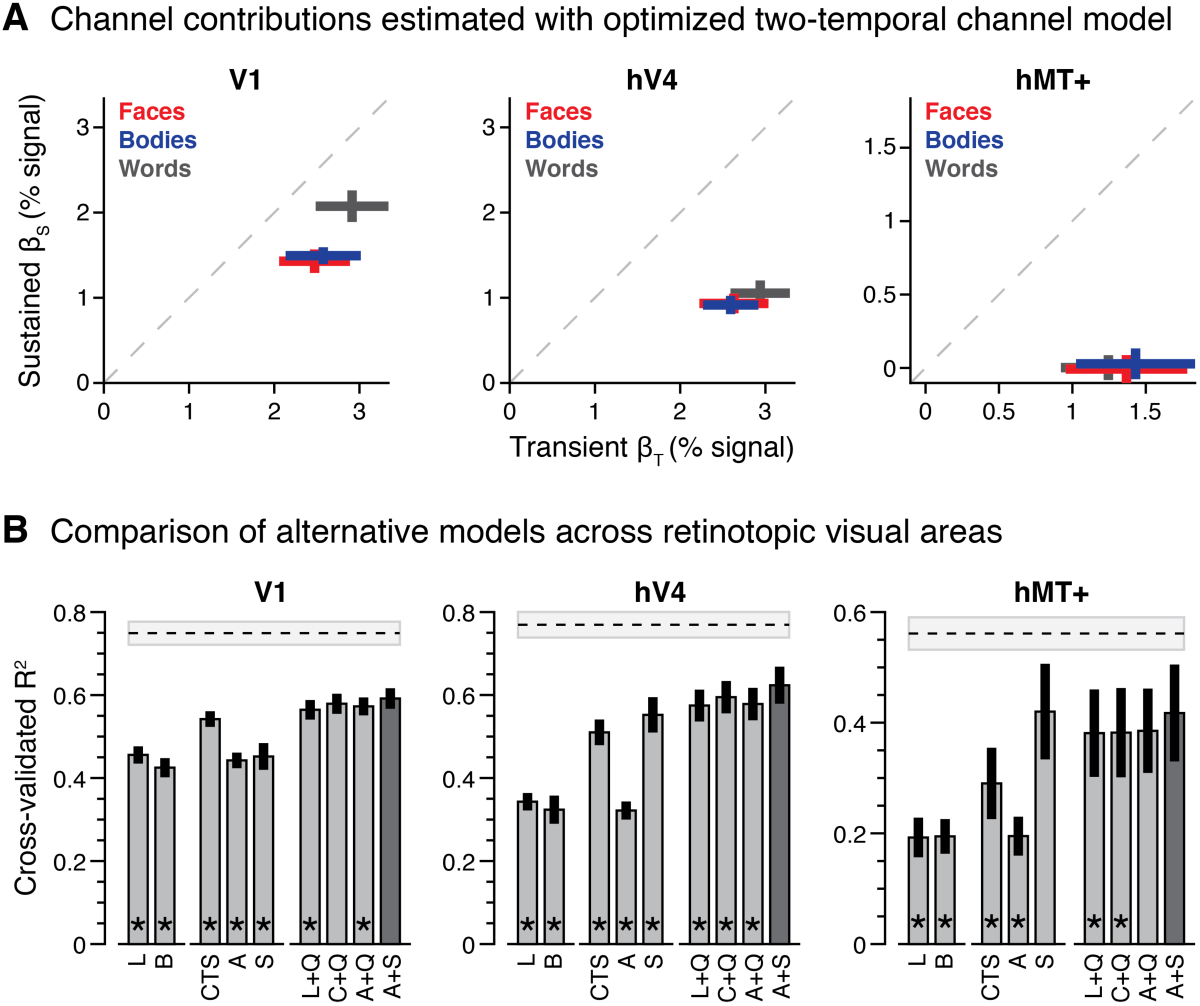
Contributions of transient and sustained temporal channels across early visual areas. (*A*) Contributions (*β* weights) of transient (*x* axis) and sustained (*y* axis) channels for each stimulus category estimated by the two-temporal channel A+S model in V1, hV4, and hMT+. Crosses span ±1 SEM across participants in each axis, and *β* were solved by fitting the model using data concatenated across all experiments. Data show average model weights across all splits of the data for each participant. *Red:* response to faces. *Blue:* response to bodies. *Gray:* response to words. *Dashed gray*: identity line (*β*_S_ = *β*_T_). (*B*) Comparison of model performance (cross-validated *R*^2^) in each region averaged across all three experiments. Hemodynamic models: *L* and *B*. Single-channel neural models: *CTS*, *A*, and *S*. Two-channel neural models: *L+Q* (5), *C+Q*, *A+Q, and A+S*. Asterisks denote models with significantly different performance vs. the A+S model.

**Figure S5.**
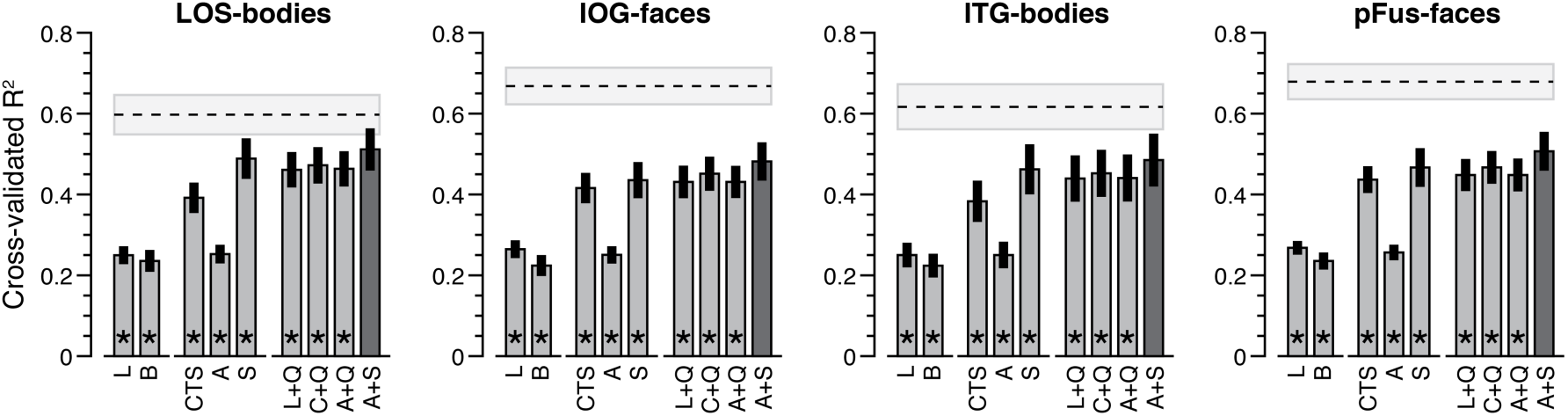
Comparison of temporal encoding models across occipital and posterior ventral temporal visual cortex. Comparison of model performance (cross-validated *R*^2^) in each region averaged across all three experiments. Hemodynamic models: *L* and *B*. Single-channel neural models: *CTS*, *A*, and *S*. Two-channel neural models: *L+Q* (5), *C+Q*, *A+Q, and A+S*. Asterisks denote models with significantly different performance vs. the A+S model.

